# Remodeling of organelles and microtubules during spermiogenesis in the liverwort *Marchantia polymorpha*

**DOI:** 10.1101/2021.07.10.451882

**Authors:** Naoki Minamino, Takuya Norizuki, Shoji Mano, Kazuo Ebine, Takashi Ueda

## Abstract

Gametogenesis is an essential biological event for sexual reproduction in various organisms. Bryophytes and some other plants employ motile sperms (spermatozoids) as male gametes, which self-locomote to the egg cells to accomplish fertilization. Spermatozoids of bryophytes harbor distinctive morphological characteristics, including the cell body with a helical slender shape and two motile flagella at the anterior edge. During transformation from a spermatid to spermatozoid (spermiogenesis), the shape and cellular contents of spermatids are dynamically reorganized. However, how each organelle is reorganized during plant spermiogenesis remains obscure. In this study, we classified the developmental processes during spermiogenesis in the liverwort *Marchantia polymorpha* according to cellular and nuclear shapes and flagella development. We then examined the remodeling of microtubules and reorganization of endomembrane organelles during spermiogenesis. The results indicate that the state of post-translational modification of tubulin is dynamically changed during the formation of the flagella and spline, and the plasma membrane and endomembrane organelles are drastically reorganized in a precisely regulated manner during spermiogenesis. These findings are expected to provide useful indexes to classify developmental and subcellular processes of spermiogenesis in bryophytes.

**Summary statement:** We classified developmental processes of spermatozoids into 1 + 5 stages and characterized remodeling of organelles and microtubules during spermiogenesis in the liverwort *Marchantia polymorpha*.

## Introduction

Sexual reproduction, one of the most crucial events in the lifecycle of multicellular organisms, is necessary for genetic variation and thus adaptation to environments. To accomplish sexual reproduction, reproductive cells, which are generally classified into male and female gametes, are generated through precisely regulated differentiation and maturation processes. A majority of seed plants generate non-motile male gametes, which are transported to the egg cells via pollen tubes for fertilization. Conversely, some species in charophycean algae as well as bryophytes, lycophytes, ferns, and some groups of gymnosperms, such as, Ginkgo and cycads, generate motile sperm cells (spermatozoids) as male gametes, which harbor two or more motile flagella and move to egg cells in water to accomplish fertilization. Gametogenesis in angiosperms has been intensively studied using several species including Arabidopsis, and its mechanisms are well documented at the genetic and molecular levels (Berger and Twell, 2011; Hackenberg and Twell, 2019; Russell and Jones, 2015; Twell, 2011). Conversely, molecular genetic studies of gametogenesis in basal land plants producing spermatozoids have only recently begun, using a few bryophyte models such as *Physcomitrium patens* and *Marchantia polymorpha* (Higo et al., 2018; Koi et al., 2016; Koshimizu et al., 2018; Minamino et al., 2017; Sanchez-Vera et al., 2017). Spermatozoids of bryophytes have distinctive morphological characteristics, some of which are also shared with spermatozoids of other plant lineages. The cell body exhibits a helical shape and consists mostly of an elongated nucleus. The anterior region of the cell body contains one mitochondrion, which is associated with the multilayered structure (MLS), a characteristic of plant spermatids. The uppermost layer of the MLS (the opposite side to the mitochondrion) is a bundle of microtubules, termed the spline, which extends along the helical nucleus through the cell body. Two flagella elongate from two basal bodies, which are attached to the spline of the MLS. The posterior side of the cell body contains the other mitochondrion and a plastid, and other cytoplasmic components are not appreciably observed in mature spermatozoids (Fig. 1A; Shimamura, 2016). During the transformation from spermatids to spermatozoids, which is termed spermiogenesis, the cell shape and cellular contents of spermatids are drastically reorganized. The MLS and flagella are formed *de novo* during spermiogenesis, the process that has been investigated mainly using electron microscopy (Carothers and Kreitner, 1967; Carothers and Kreitner, 1968; Kreitner, 1977a; Kreitner and Carothers, 1976; Renzaglia and Duckett, 1987; Renzaglia and Garbary, 2001). It is expected that most of the cytoplasm, including various organelles other than the nucleus, anterior and posterior mitochondria, and a plastid, is eliminated during spermiogenesis. However, when and how organelles are removed from spermatids during spermiogenesis remains unknown, mainly due to the lack of proper methods to observe spatiotemporal organelle dynamics during plant spermiogenesis.

**Figure 1.**
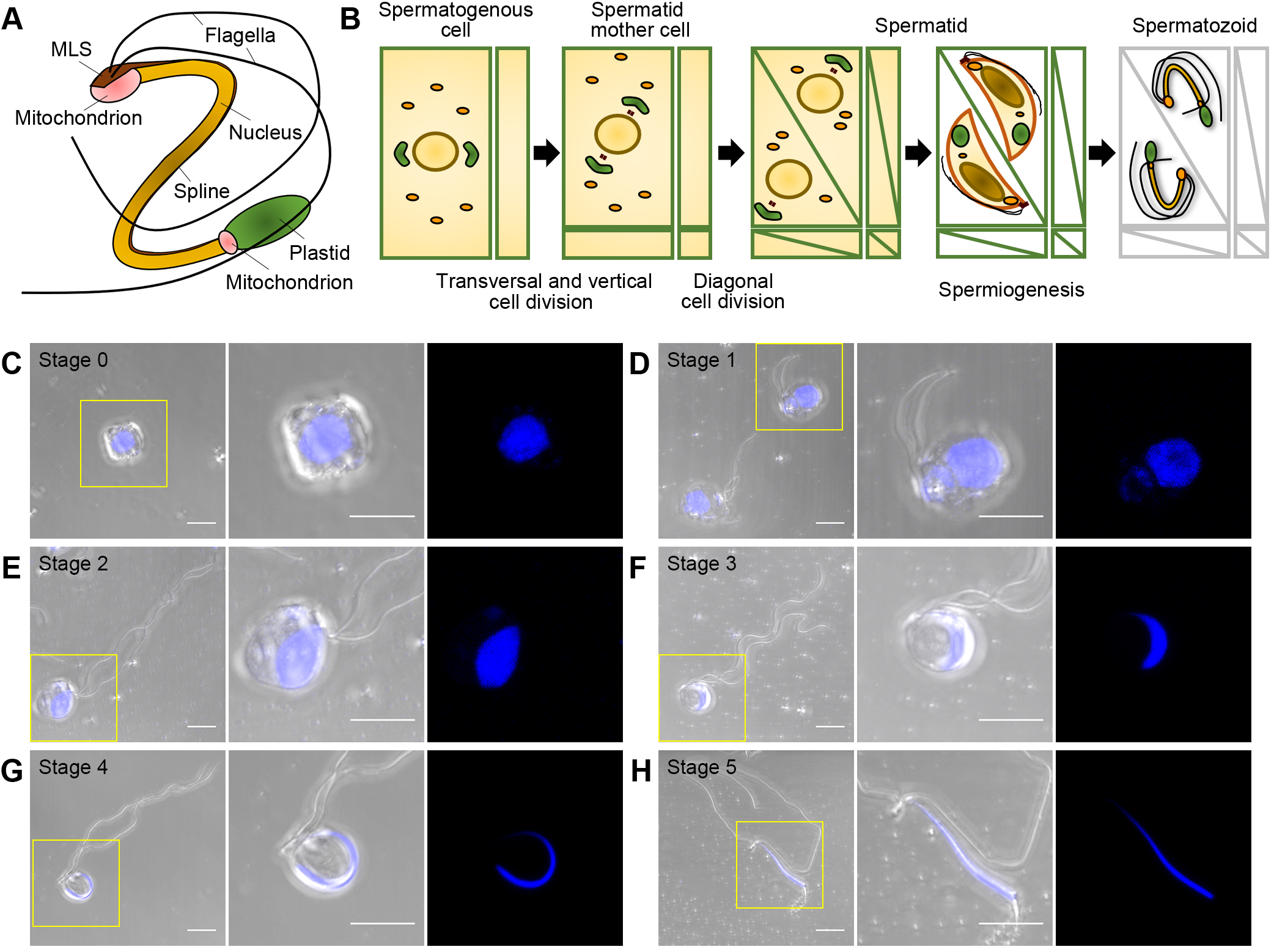
Developmental stages of spermiogenesis in *M. polymorpha*. (A-B) Schematic illustration of spermatozoids (A) and male gamete development (B). (C-H) Maximum intensity projection images of spermatids and a spermatozoid stained with Hoechst 33342. Left panels, merged images of differential interference contrast (DIC) and fluorescent images; middle panels, higher magnification images of left panels; right panels, fluorescent images of nuclei in middle panels. The blue pseudo-color indicates fluorescence from Hoechst 33342. Scale bars = 5 μm.

*Marchantia polymorpha* is a model of liverworts, a group of bryophytes whose genomic information is available (Bowman et al., 2017; Montgomery et al., 2020), and various tools for molecular genetic analyses have been established (Kohchi et al., 2021). Transcriptome data and a wide variety of organelle markers have also been developed for *M. polymorpha* (Bowman et al., 2017; Higo et al., 2016; Kanazawa et al., 2016; Minamino et al., 2017; Minamino et al., 2018). Employing this relatively new model plant for analyzing the cellular dynamics during plant spermiogenesis, we previously reported that subcellular localizations of some organelle markers drastically change in spermatogenous tissue (Minamino et al., 2017). However, the temporal resolution was not sufficient because of the lack of defined developmental stages in spermiogenesis in *M. polymorpha*. In this study, we first classified the developmental processes of spermiogenesis in *M. polymorpha* according to cellular and nuclear morphology and flagellar formation. We then examined microtubule organization in each developmental stage during spermiogenesis and found that the state of post-translational tubulin modification is significantly changed during the development of microtubule-containing structures. Furthermore, we monitored markers of endomembrane organelles in each developmental stage and found that each organelle/organelle protein is degraded at a distinctive stage during spermiogenesis. These results demonstrate a drastic but highly organized rearrangement of cellular components during spermiogenesis in *M. polymorpha* and provide a unified standard for classification of developmental stages of spermatids undergoing spermiogenesis in bryophytes.

## Results

### Determining developmental stages of spermiogenesis in *M. polymorpha*

An antheridium of *M. polymorpha* consists of outer jacket cells and inner reproductive cells. Spermatogenous cells proliferate by continuous transverse and vertical cell division, and the resultant spermatid mother cells divide diagonally to generate spermatids (Fig. 1B; Shimamura, 2016). Spermatids then transform into mature spermatozoids through spermiogenesis. Spermiogenesis of *M. polymorpha* comprises dynamic cellular reorganization events, including condensation and elongation of the nucleus, *de novo* synthesis of the locomotory apparatus, and elimination of the cytoplasm (Fig. 1B; Shimamura, 2016). Based on the morphology of the cell body, nuclear shape observed with Hoechst 33342, and formation of flagella as yardsticks, we first attempted to classify spermatid cells undergoing spermiogenesis, which were fixed with paraformaldehyde and separated from each other by treating with cell-wall digesting enzymes, into 1 + 5 developmental stages (Fig. 1C-H). The cells harboring the round nucleus without visible flagella were classified as stage 0. These cells presumably contained spermatogenous cells, spermatid mother cells, and early-stage spermatids, which were not distinguishable from each other based on their morphology after cell wall digestion (Fig. 1C). At stage 1, protruding or elongating flagella were observed, whereas the nucleus remained spherical (Fig. 1D). A change in the nuclear shape was detected beginning from stage 2. At this stage, spermatids were equipped with fully elongated flagella, and projection of the anterior side of the nucleus was also visible (Fig. 1E). At stage 3, the nucleus became crescent-shaped (Fig. 1F), and a cylindrical nucleus was observed at stage 4 (Fig. 1G). Mature spermatozoids were defined as stage 5 (Fig. 1H).

### Altered modification of microtubules during spermiogenesis in *M. polymorpha*

The spermatozoid of *M. polymorpha* comprises two distinctive microtubule-containing structures, the flagella and spline. To examine how microtubules are organized during spermiogenesis to form these structures, we performed immunostaining of tubulin in spermatids at distinct developmental stages. In addition to immunostaining using an anti-α-tubulin antibody, we also performed staining with an anti-poly-glutamate (polyE) antibody, which recognizes the carboxyl-terminally located linear alpha-glutamate chain comprising four or more glutamate residues. This method was chosen knowing that post-translational modifications, including glutamylation, are detected in persistent microtubules in the axoneme in animals and in Chlamydomonas (Janke and Magiera, 2020; Wloga et al., 2017). In some cells at stage 0, only the anti-α-tubulin antibody-stained fibrous structures were observed near the cell surface (Fig. 2A). In other cells, rod-like forming flagella, which had not yet protruded from the cell body, were stained by both the anti-α-tubulin and anti-polyE antibodies (Fig. 2B). In stage 1 spermatids, elongating flagella were observed, stained by both antibodies (Fig. 2C). Notably, microtubule fibers extending radially from or toward the basal region of the forming flagella were frequently observed at stages 0 and 1 (Fig. 2B,C). The radial fibers were not observed in stage 2 spermatids, whereas the spline microtubule was stained by the anti-α-tubulin antibody in stage 2 and 3 spermatids (Fig. 2D,E). The radial structure and spline were not stained by polyE antibody (Fig. 2B-E). Intriguingly, the signal intensity from α-tubulin on flagella became weaker except for that from the distal-most region at stages 2 to 3; however, the signal from polyE was uniformly detected at the corresponding region (Fig. 2D,E). Similar results were also obtained when we used another anti-α-tubulin and anti-β-tubulin antibodies (Fig. S2A-D). In stage 4, the polyE signal was detected on flagella, but α-tubulin was not detected on either the flagella or spline (Fig. 2F). At stage 5 (mature spermatozoids), the α-tubulin signal was detected again on both flagella and spline. The polyE signal was also detected on flagella and spline, although the intensity of the signal from the spline was weak compared with that from the flagella (Fig. 2G; Fig. S1). We then performed immunoblotting to examine whether the anti-polyE antibody recognized the same tubulin population as anti-α- and anti-β-tubulin antibodies. The anti-α- and β-tubulin antibodies detected tubulin molecules at approximately 50 kDa, whereas the anti-polyE antibody reacted with the product at approximately 58 kDa in cell lysates prepared from antheridia (Fig. S3). The anti-α- and -β-tubulin antibodies did not react with the 58 kDa product detected with the anti-polyE antibody, and vice versa. Given that the shift of the molecular weight would reflect the covalently added poly-glutamate chain, as reported in previous studies (Kubo et al., 2010; Kubo et al., 2014; Vu et al., 2016), and that both the epitopes of the anti-tubulin antibodies used in this study and polyglutamylation sites are mapped to the C-terminal region of tubulin, the anti-tubulin antibodies and the anti-polyE antibody react exclusively with distinct populations of tubulin molecules. Thus, our results strongly suggest that tubulin molecules in the axoneme and spline undergo progressive alteration in post-translational modification during spermiogenesis.

**Figure 2.**
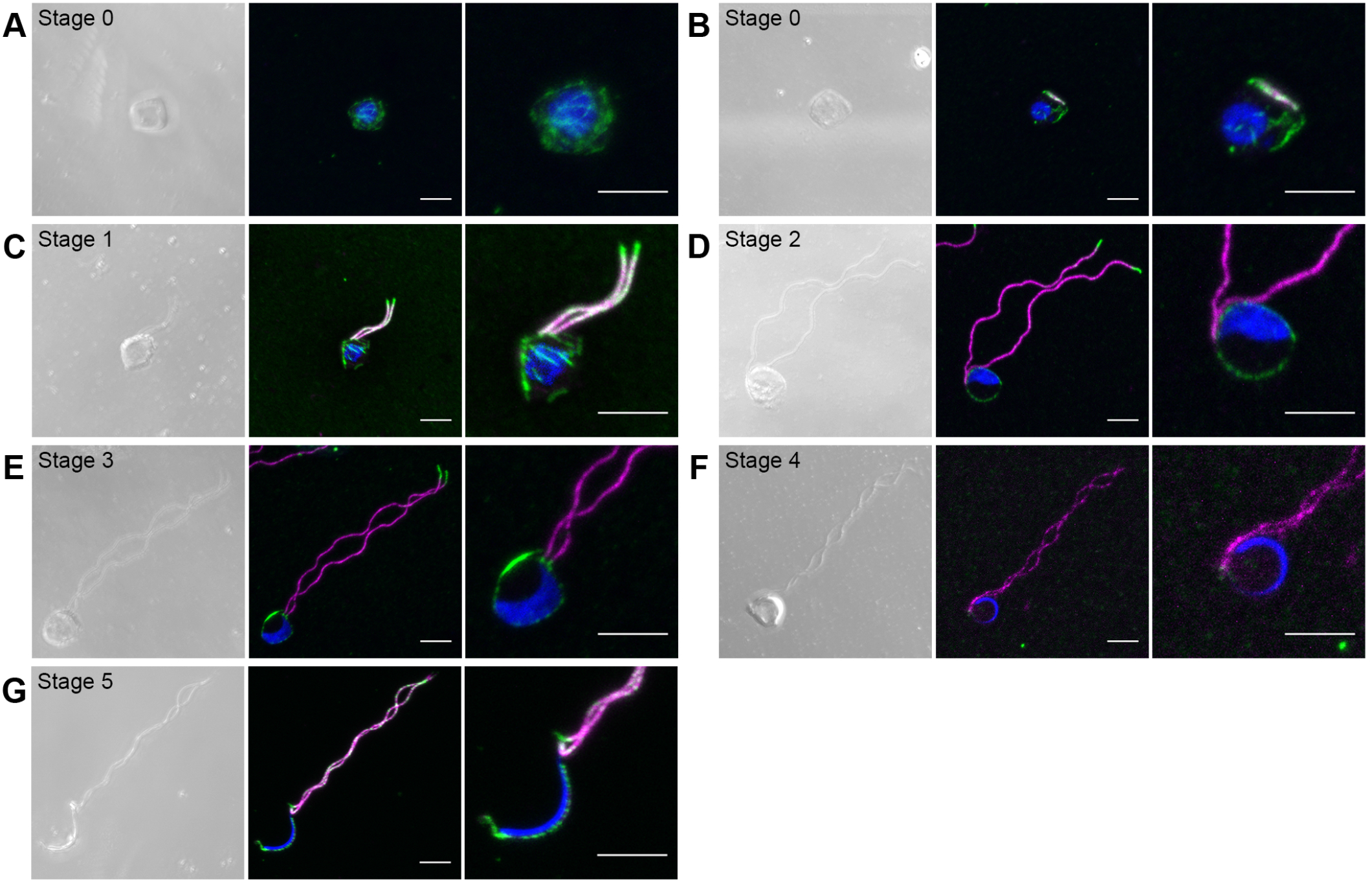
Microtubule dynamics during spermatogenesis. (A-G) Maximum intensity projection images of spermatids and a spermatozoid immunostained with anti-α-tubulin and anti-polyE antibodies. Left panels, differential interference contrast (DIC) images; middle panels, fluorescent images; right panels, higher magnification images of middle panels. Green, magenta, and blue pseudo-colors indicate fluorescence from Alexa 488 (α-tubulin), Alexa594 (polyE), and Hoechst 33342, respectively. Scale bars = 5 μm.

### Selection of promoters for expression of organelle markers in spermatogenous tissue of *M. polymorpha*

With the goal of observing the behavior of endomembrane organelles during spermiogenesis without interference from surrounding tissues, we expressed organelle markers under the regulation of promoters predominantly active in spermatids. We selected three candidate genes, which were reported to express strongly during spermiogenesis in *M. polymorpha*. Mp*DUO1* is a MYB transcription factor, which is required for sperm differentiation of *M. polymorpha* (Higo et al., 2018). Mp*CEN1* is a homolog of CENTRIN essential for the formation of locomotory apparatus in the fern *Marsilea vestita* and is strongly expressed in spermatogenous tissue independent of Mp*DUO1* in *M. polymorpha* (Higo et al., 2018; Klink and Wolniak, 2001). IFT52 is one of the components of the intraflagellar transport complex essential for flagellar formation in Chlamydomonas (Brazelton et al., 2001; Deane et al., 2001), whose homolog, Mp*IFT52*, in *M. polymorpha*, is abundantly expressed in antheridiophores according to the transcriptome data deposited in MarpolBase (https://marchantia.info/). Transgenic lines expressing the fluorescent protein Citrine driven by the promoters of these genes exhibited strong expression in spermatogenous tissue (Fig. S4A-C), suggesting that these promoters may be useful for expression of organelle markers during spermatogenesis. We thus used these promoters for further experiments in this study (Fig. S4D).

### Reorganization of vacuoles during spermiogenesis in *M. polymorpha*

We previously reported that plasma membrane (PM) proteins and several endomembrane organelle proteins are transported into the luminal space of spherical vacuoles to be degraded during spermiogenesis (Minamino et al., 2017). However, it remains unknown when the vacuole transforms to spherical shape, and it also remains unclear how transport of organelle proteins to the vacuole is organized. To determine this, we first examined alteration of the vacuolar morphology during spermiogenesis using a vacuole-residing soluble *N*-ethylmaleimide sensitive factor attachment protein receptor (SNARE), MpVAMP71, tagged with monomeric Citrine (mCitrine-MpVAMP71), expressed under the regulation of the Mp*DUO1* promoter (Kanazawa et al., 2016). In cells at stages 0 and 1, small and fragmented vacuoles with intricate morphology were observed around the nucleus (Fig. 3A,B). However, only one spherical vacuole was observed in each spermatid at stage 2 (Fig. 3C), which persisted through stages 3 and 4 (Fig. 3D,E). In mature spermatozoids (stage 5), compartments with mCitrine-MpVAMP71 were not detected (Fig. 3F). Thus, the vacuole drastically changes its morphology and number before stage 2 to form a spherical vacuole, which is eliminated after stage 4 to accomplish spermiogenesis.

**Figure 3.**
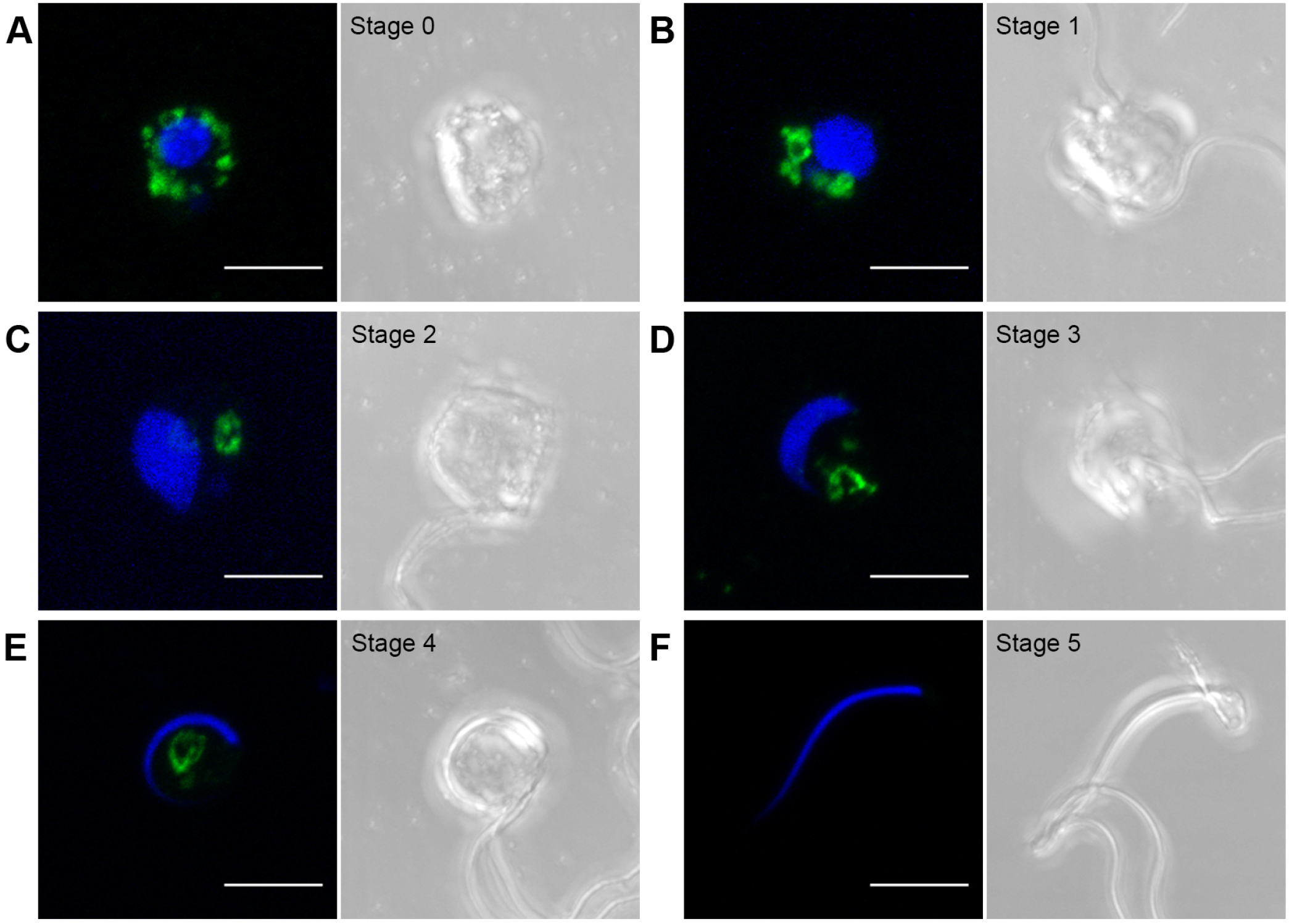
Changes in vacuolar morphology during spermiogenesis. (A-F) Maximum intensity projection images of spermatids and a spermatozoid expressing mCitrine-MpVAMP71 driven by the Mp*DUO1* promoter. Left panels, fluorescent images; right panels, differential interference contrast (DIC) images. Green and blue pseudo-colors indicate fluorescence from mCitrine and Hoechst 33342, respectively. Scale bars = 5 μm.

### Dynamic reorganization of the plasma and endo-membranes during spermiogenesis

A plasma membrane SNARE, MpSYP12A (Kanazawa et al., 2016), tagged with mCitrine (mCitrine-MpSYP12A), was shown to be removed from the PM during spermiogenesis, suggesting that drastic remodeling also takes place in the PM in spermatids undergoing spermiogenesis (Minamino et al., 2017). To elucidate the precise timing of this event, we observed spermatids expressing mCitrine-MpSYP12A at each developmental stage. At stage 0, mCitrine-MpSYP12A was detected on the PM and punctate compartments (Fig. 4A,B). At stage 1, the fluorescent signal on the PM was diminished, and bright fluorescence was observed in punctate compartments (Fig. 4C), which probably represents small vacuoles observed in spermatids at stage 1 (Fig. 3B). The signal from mCitrine was still detectable in the lumen of the larger spherical vacuole in some spermatids at stage 2 (Fig. 4D), whereas only faint or almost no fluorescence was detected in the majority of spermatids at this stage (Fig. 4E). We did not detect any fluorescence after stage 3 (Fig. 4F-H). These results indicate that endocytosis was highly activated between stages 0 and 1, and at least a set of PM proteins including MpSYP12A was removed almost completely from the PM and degraded in the vacuole during stage 2, suggesting rapid remodeling of the PM at early stages during spermiogenesis.

**Figure 4.**
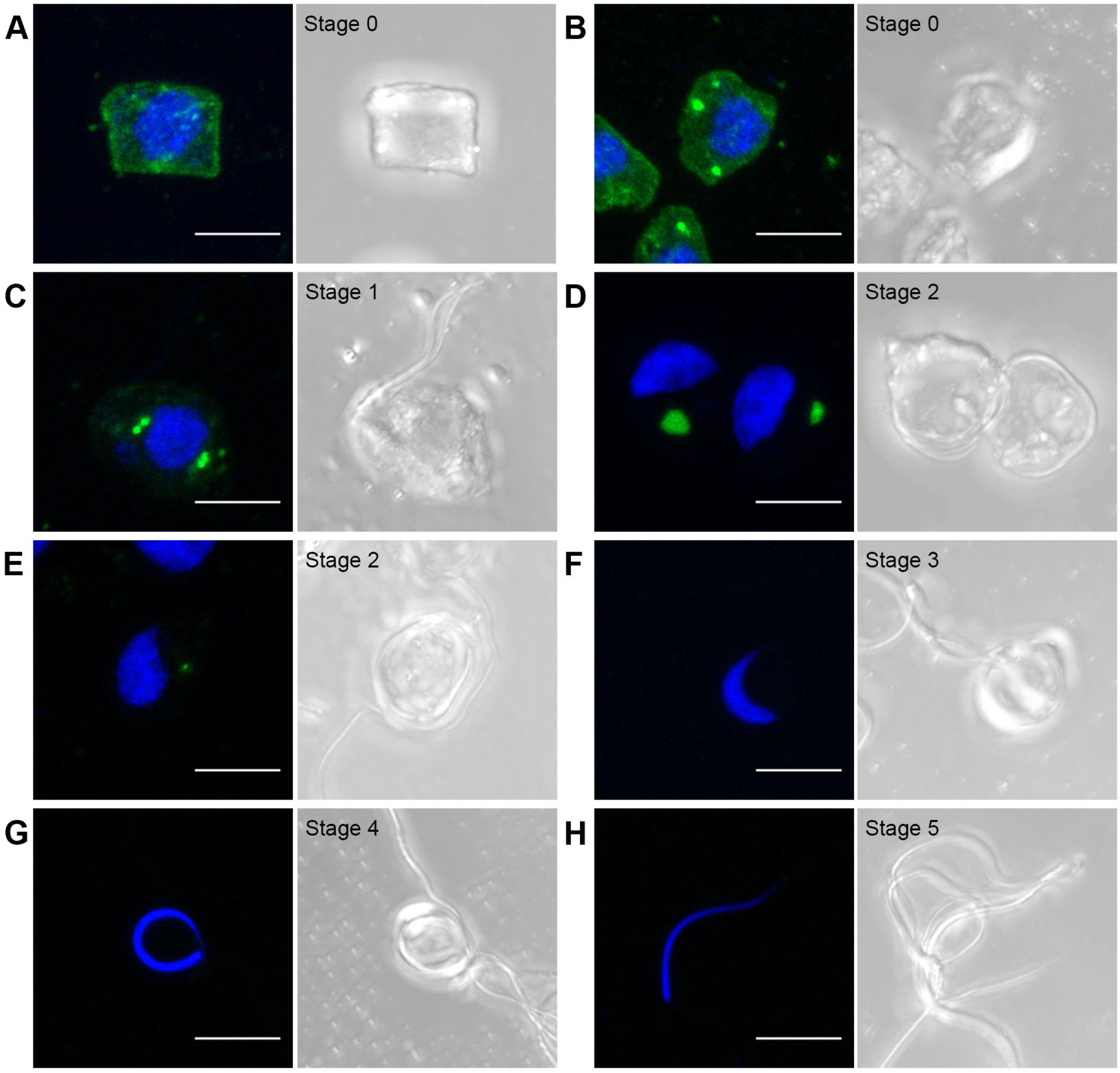
Remodeling of the plasma membrane during spermiogenesis. (A-H) Maximum intensity projection images of spermatids and a spermatozoid expressing mCitrine-MpSYP12A driven by the Mp*DUO1* promoter. Left panels, fluorescence images; right panels, differential interference contrast (DIC) images. Green and blue pseudo-colors indicate fluorescence from mCitrine and Hoechst 33342, respectively. Scale bars = 5 μm.

The Golgi apparatus also exhibited drastic changes in morphology and number during spermiogenesis. We observed the Golgi apparatus during spermiogenesis using the Venus-fused transmembrane domain of sialyltransferase (ST) derived from rat, which is localized to the Golgi apparatus in *M. polymorpha* (Kanazawa et al., 2016). While several discrete Golgi apparatuses were observed at stage 0, only one Golgi apparatus, which was enlarged compared with the Golgi at earlier stages, was observed in cells at stage 2 (Fig. 5A-C,G,H). Appearance of the large Golgi apparatus in spermatids undergoing spermiogenesis is also reported in electron microscopic observations of another liverwort, *Blasia pusilla* (Renzaglia and Duckett, 1987). The fluorescence from ST-Venus was decreased in spermatids at stage 3 and was not detectable in cells at stages 4 and 5 (Fig. 5D-F). We also observed similar behavior of the Golgi apparatus using other Golgi markers, mCitrine-fused MpGOS11, and MpSFT1 (Kanazawa et al., 2016); these markers were hardly detected at as early as stage 3 (Fig. S5). Thus, one large Golgi apparatus is formed during early stages of spermiogenesis, which is degraded during later stages with distinct temporal regulation depending on proteins. We then observed changes in the endoplasmic reticulum (ER). As an ER membrane marker, we used mCitrine-fused MpSEC22 (mCitrine-MpSEC22, Kanazawa et al., 2016). We also monitored a soluble ER protein marker SP-mCitrine-HDEL, which consists of mCitrine tagged with the signal peptide (SP) at its N-terminus and four amino acids (His-Asp-Glu-Leu) sufficient for ER localization at the C-terminus (Mano et al., 2018). These ER markers illuminated the nuclear envelope (NE) and the ER until stage 2 (Fig. 6A-F). Intriguingly, the fluorescence at the NE was hardly detected at stage 3, whereas the signal on the ER remained detectable at this stage (Fig. 6G-J), suggesting distinct regulation of remodeling between the NE and ER during spermiogenesis. Furthermore, we found that these two ER markers behaved distinctly at later stages; SP-mCitrine-HDEL remained at the anterior region of spermatozoids, and a faint signal was also detected in other regions (Fig. 6L), whereas the signal from mCitrine-MpSEC22 was not detected in spermatozoids (Fig. 6K). Thus, remodeling of the ER also takes place in a highly organized manner, with distinct temporal regulation depending on domains and proteins.

**Figure 5.**
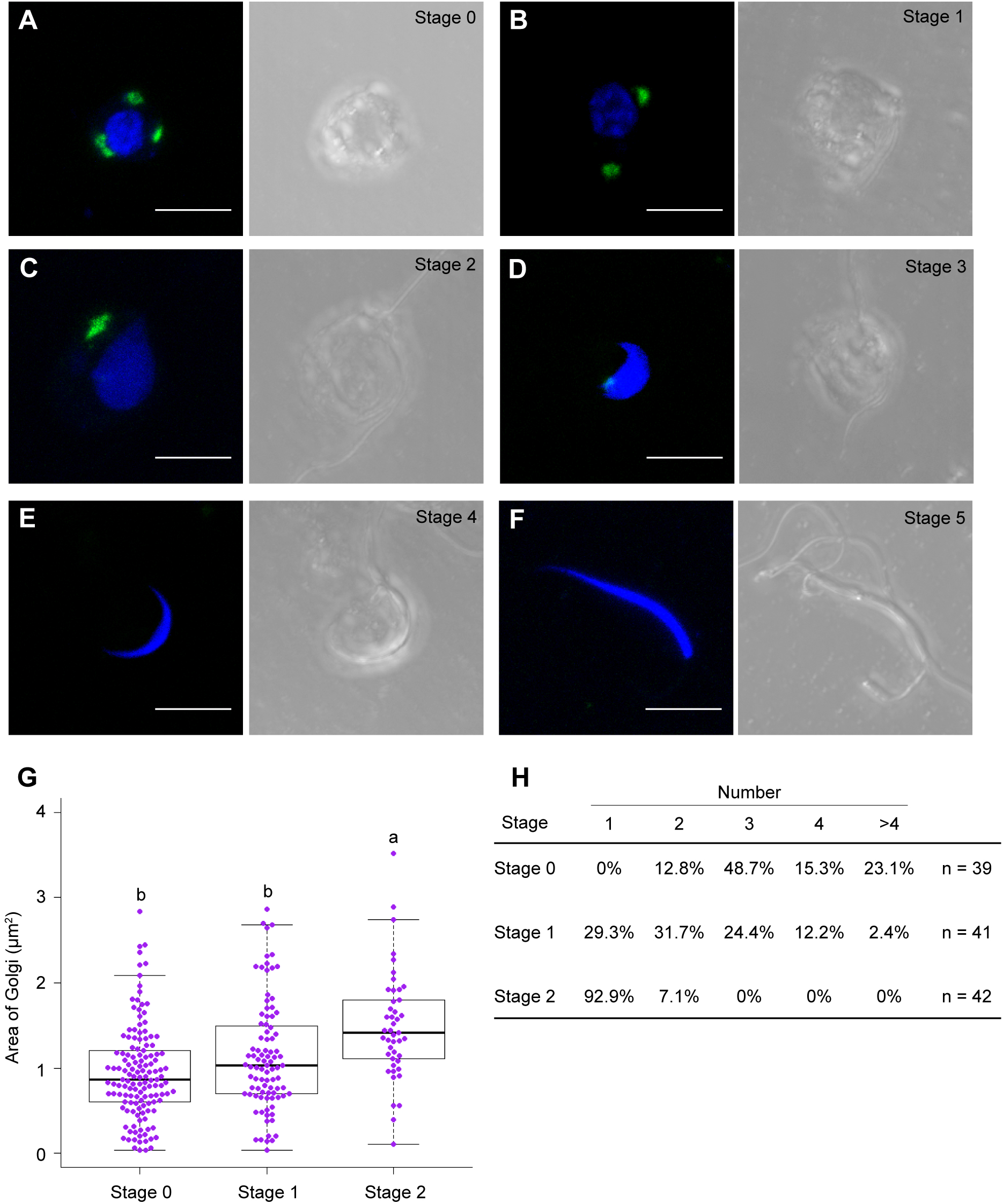
Reorganization of the Golgi apparatus during spermiogenesis. (A-F) Maximum intensity projection images of spermatids and a spermatozoid expressing ST-Venus driven by the Mp*DUO1* promoter. Left panels, fluorescence images; right panels, differential interference contrast (DIC) images. Green and blue pseudo-colors indicate fluorescence from mCitrine and Hoechst 33342, respectively. Scale bars = 5 μm. (G) The size of the Golgi apparatus labeled by ST-Venus. n = 137 (stage 0), 91 (stage 1), and 45 puncta (stage 2). The boxes and solid lines in the boxes indicate the first quartile and third quartile, and median values, respectively. The whiskers indicate 1.5 × interquartile ranges. Different letters denote significant differences based on Tukey’s test (*p* < 0.05). (H) The number of the Golgi apparatus labeled by ST-Venus. The difference in the number of the Golgi apparatus among stages was statistically significant based on Tukey’s test (*p* < 0.05). See also Fig. S5.

**Figure 6.**
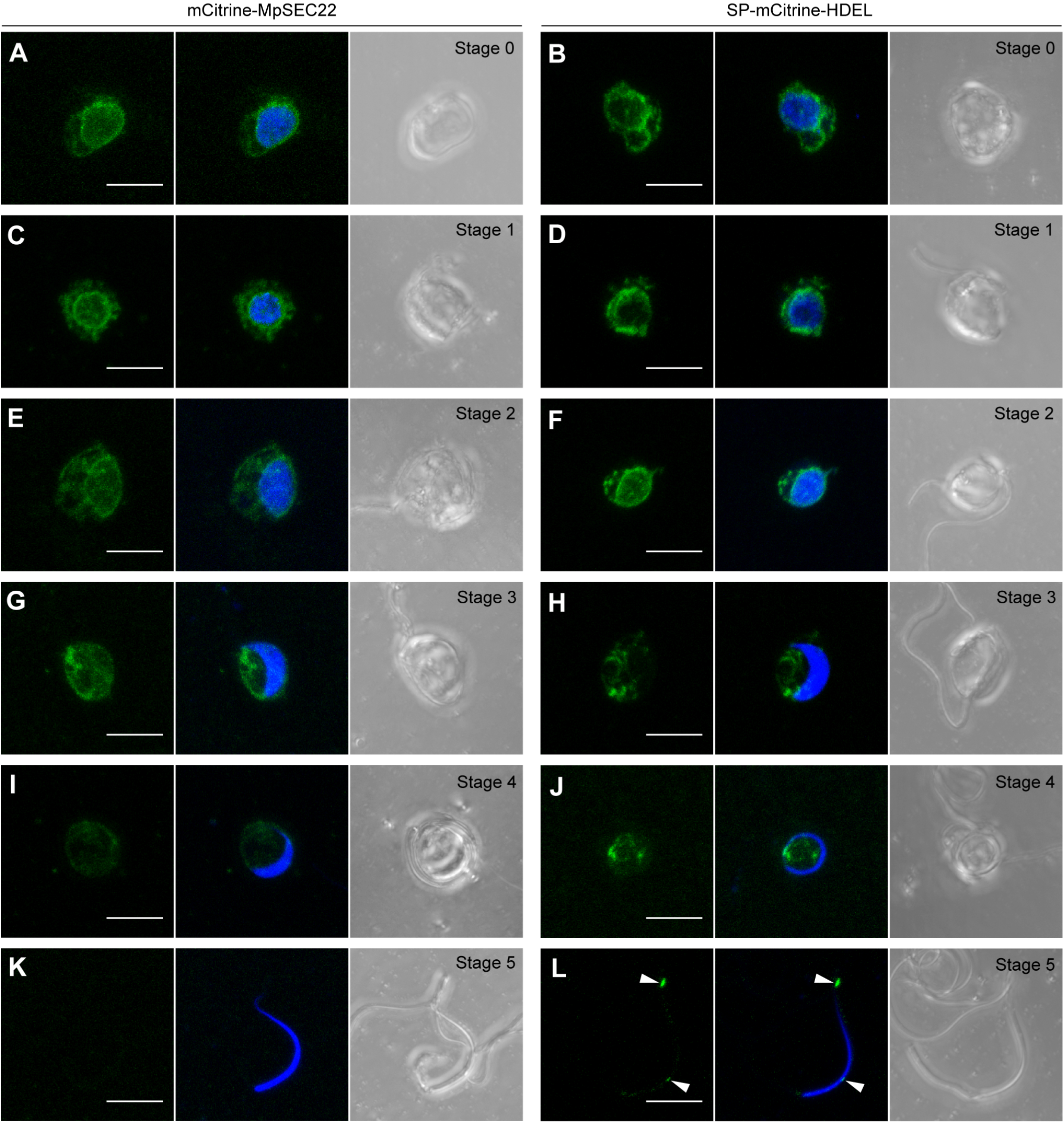
Reorganization of the ER and the nuclear envelope during spermiogenesis. Maximum intensity projection images of spermatids and spermatozoids expressing mCitrine-MpSEC22 driven by the Mp*DUO1* promoter (A, C, E, G, I, K) or SP-mCitrine-HDEL driven by the Mp*EF1*α promoter (B, D, F, H, J, L). Arrowheads indicate SP-mCitrine-HDEL remaining in spermatozoids. Left panels, fluorescence images of mCitrine; middle panel, merged images of mCitrine and Hoechst 33342; right panels, differential interference contrast (DIC) images. Green and blue pseudo-colors indicate fluorescence from mCitrine and Hoechst 33342, respectively. Scale bars = 5 μm.

## Discussion

The unified criterion for staging developmental processes is useful for developmental studies of any organisms, as exemplified by human embryogenesis or spermatogenesis staging, or development of the flower or anther in Arabidopsis (Bowman et al., 1991; Meistrich and Hess, 2013; O’Rahilly and Müller, 1987; Oakberg, 1956; Sanders et al., 1999). A shared criterion for staging developmental processes would also be quite useful in studies of plant spermatozoid development. In this study, we propose 1 + 5 stages of spermiogenesis in *M. polymorpha*, according to the shape of the cell body and nucleus and flagellar formation (Fig. 7). The continuous morphological changes of the nucleus in fixed spermatids were roughly classified as spherical, drop-shaped, crescent-shaped, or cylinder-shaped (Fig. 1A-F), which is also consistent with previous observations using electron microscopy (Kreitner, 1977b). The change in nuclear morphology during spermiogenesis is accompanied by chromatin condensation, which is associated with conversion of DNA-binding nuclear basic proteins from histones to protamine-like proteins (D’Ippolito et al., 2019; Higo et al., 2016). Chromatin condensation also occurs during spermiogenesis in animal systems, and transcription ceases completely around mid-spermiogenesis as chromatin condensation progresses (Kierszenbaum and Tres, 1975; Sassone-Corsi, 2002). Further progression of spermiogenesis is mediated by translating mRNA that is accumulated before spermiogenesis at appropriate time points (Chalmel and Rolland, 2015; Dai et al., 2019). Similar cessation of transcription may occur during spermatogenesis in plants; therefore, morphological characteristics, rather than transcriptome information, would be useful indexes for distinguishing developmental stages of plant spermiogenesis.

**Figure 7.**
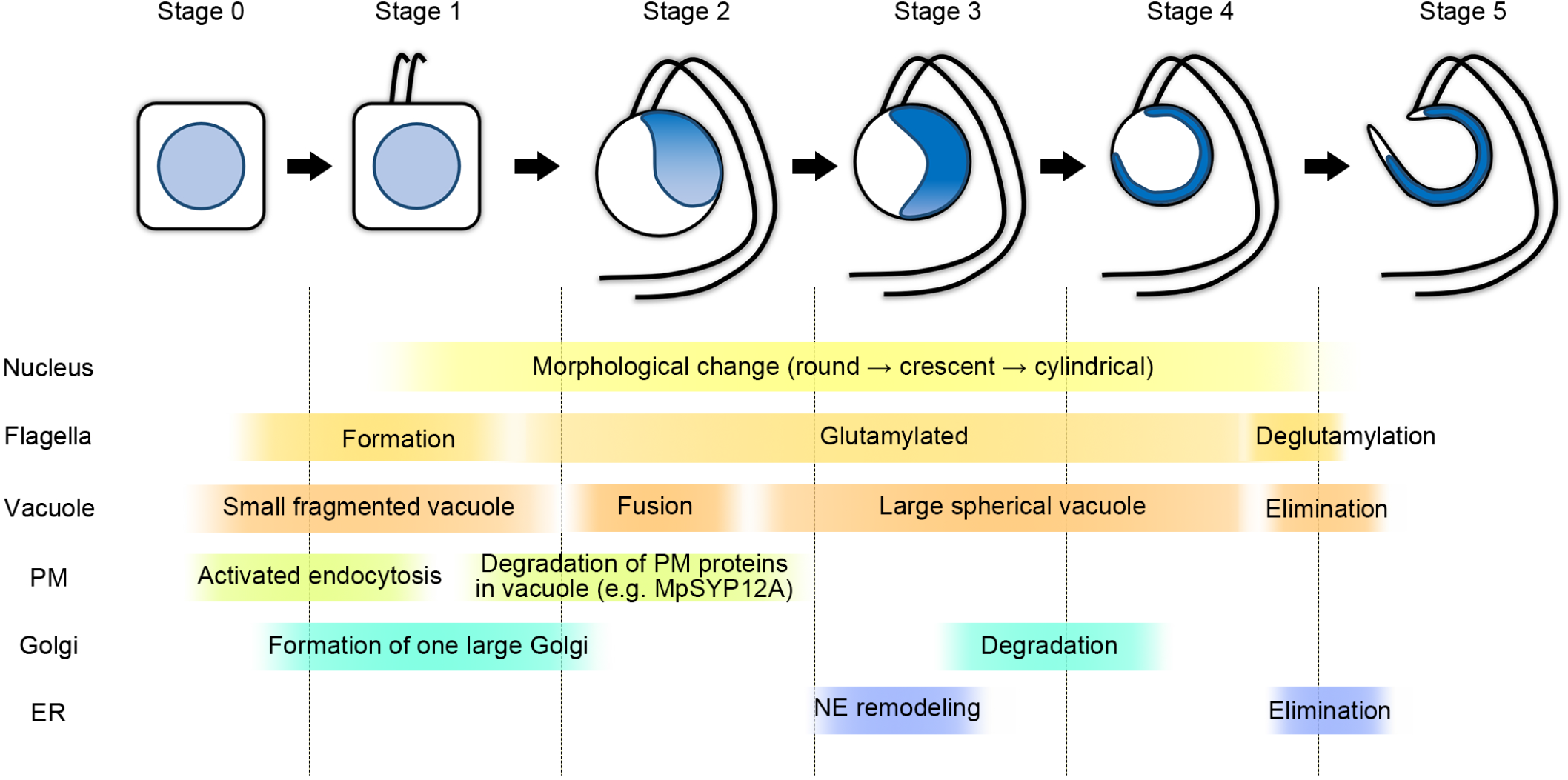
Schematic diagram of spermiogenesis progression in *M. polymorpha*. At stage 0, flagellar formation begins in a cell body, and endocytosis is highly activated. At stage 1, flagella protrude and elongate. Radially extending microtubules are formed transiently from or to the basal region of the flagella. Plasma membrane (PM) proteins including mCitrine-MpSYP12A are transported to vacuoles. At stage 2, flagella elongation is mostly completed, and morphological change of the nucleus begins. Spline microtubules are visible. Vacuoles fuse with each other to form a large spherical vacuole. PM proteins are degraded in the spherical vacuoles. One large Golgi apparatus is observed. At stage 3, Golgi proteins are degraded. The nuclear envelope is remodeled. At stage 4, most of endoplasmic reticulum (ER) proteins are eliminated. The spherical vacuole persists until stage 4, which is removed to transform to the mature spermatozoid. Glutamylation of tubulin in axonemal and spline microtubules is dynamically regulated during spermiogenesis.

After dividing the process of transformation from spermatids to spermatozoids into 1 + 5 stages, we analyzed microtubule organization in spermatids undergoing spermiogenesis and spermatozoids by immunostaining with antibodies for tubulin or a post-translational modification of tubulin (polyE). At stage 1, microtubules radially extending from the basal region of elongating flagella were observed (Fig. 2B,C). This structure was a transient structure and disappeared before stage 2, at which point flagellar elongation seemed to be nearly complete (Fig. 2D). Therefore, the radial microtubules might function as rails for transporting components of flagella to the bases of the flagella. A similar structure, termed rootlet microtubules, has been observed in Chlamydomonas and which extends from the basal bodies and have been involved in arrangement of the eyespot (Boyd et al., 2011). Although spermatids of *M. polymorpha* do not contain the eye spot, the radial structure might be involved in the arrangement of organelles, such as mitochondria, during spermatogenesis. Tubulin of the basal bodies and axonemes undergoes several post-translational modifications including glutamylation (Janke and Magiera, 2020; Wloga et al., 2017). We found that the flagella and spline in spermatozoids of *M. polymorpha* were recognized by the anti-polyE antibody (Fig. 2; Fig. S1, S2), indicating that axonemal and spline microtubules are glutamylated in the basal land plant. Glutamylation is reported to be involved in regulation of flagellar motility in Chlamydomonas and Tetrahymena (Kubo et al., 2010; Suryavanshi et al., 2010), assembly of doublet microtubules in mouse (Konno et al., 2016; Lee et al., 2013), and stability of microtubules in *Caenorhabditis elegans* (O’Hagan and Barr, 2012; O’Hagan et al., 2011). The function of the glutamylation of axonemal and spline microtubules in plant spermatozoids should be verified in future studies.

We found that the signals of anti-tubulin antibodies used for immunostaining in this study became weaker as spermiogenesis progressed and then recovered in mature spermatozoids (Fig. 2; Fig. S2). One plausible explanation of this phenomenon is that the status of post-translational modification of tubulin, including glutamylation, changes as spermiogenesis progresses. The α- and β-tubulin molecules are poly-glutamylated at the C-terminal region (Wloga et al., 2017), and epitopes of anti-tubulin antibodies used in this study are also located in the C-terminal region of tubulin molecules. Thus, glutamylation could hinder access of the anti-tubulin antibodies to their epitopes, resulting in diminished signals in immunostaining. This is consistent with the result of immunoblotting; anti-α- and -β-tubulin antibodies did not recognize poly-glutamylated tubulin, which was detected by the anti-polyE antibody (Fig. S3). Thus, our results indicate that an extent of glutamylation of tubulin molecules may change during spermiogenesis; the degree of glutamylation increases along formation of the axoneme, and after its completion, microtubules might be de-glutamylated. An effect of post-translational modification of tubulin on microtubule properties in spermatids and spermatozoids and its physiological significance would be interesting topics for further study.

We investigated the reorganization of endomembrane organelles with the progression of spermiogenesis; the results are summarized in Figure 7. The vacuole changed its morphology and number from a complicated and fragmented shape to a spherical structure at stage 3, when the nucleus assumed a crescent-like shape (Fig. 3C,D). This morphological transition suggests that vacuoles actively fuse with each other as spermiogenesis proceeds. It would be interesting to see whether evolutionarily conserved machinery components of vacuole biogenesis, such as RAB7/RABG GTPase and the HOPS complex (Brillada et al., 2018; Rojo et al., 2001; Takemoto et al., 2018), are involved in vacuole remodeling during spermiogenesis in *M. polymorpha*, which would also provide useful information for understanding its biological significance. Stage 4 spermatids contained one spherical vacuole in the cell body, whereas the mature spermatozoid had no discernable vacuoles or remnants of the vacuole (Fig. 3E,F). Thus, it appears that the vacuole is removed from the spermatid at the final step of spermiogenesis. In spermiogenesis of Drosophila, the individualization complex comprising actin cones is required for elimination of unneeded organelles and the cytosol. The complex is initially formed near the nucleus and moves from the head to the tail of the spermatid. During this movement, majority of the cytoplasm is removed to form the cystic bulge, which is finally detached from the tip of the tail (Fabian and Brill, 2012). Similarly in plants, actin seems to play an important role in the cytoplasm removal during spermiogenesis, although a structure corresponding to the individual complex has not been reported (Renzaglia and Garbary, 2001).

Our observation of mCitrine-MpSYP12A revealed that endocytosis of this protein is highly active between stages 0 and 1 (Fig. 4). This result suggests that the protein composition of the PM drastically changes at the early stage of spermiogenesis. The spermatozoid PM must have competence that is distinct from that of the spermatid, such as the ability to respond to an attractant from egg cells and resistance to low osmolarity to prevent rupture in fresh water without the rigid cell wall. Therefore, the cell surface of the spermatozoid is reorganized during spermiogenesis, and the high endocytic activity observed in early spermiogenesis might contribute to the thorough remodeling of the PM.

The morphology of the Golgi apparatus has diverged among organisms and even among tissues, probably reflecting diverse and tissue-specific Golgi functions (Ito et al., 2014; Sengupta and Linstedt, 2011). We found that the number and size of the Golgi apparatus changed during spermiogenesis (Fig. 5, Fig. S5), which suggests that the function of the Golgi apparatus changes during spermiogenesis. Furthermore, we found that the signal from Golgi markers disappeared by stage 4, which strongly suggests that the Golgi apparatus (or a major set of Golgi proteins) is eliminated before removal of the cytoplasm. Although the molecular mechanism of the Golgi removal remains ambiguous, we previously found that autophagy is highly activated during spermiogenesis and a Golgi-like membrane structure is transported into the vacuole in *M. polymorpha* (Minamino et al., 2017). Therefore, autophagy could be a key mechanism for Golgi clearance during spermiogenesis, which would be verified in future studies.

Drastic deformation of the nucleus during spermiogenesis is generally observed in land plants (Renzaglia and Garbary, 2001). However, the mechanisms of the nuclear remodeling remain unknown. We found that the signal from SP-mCitrine-HDEL and mCitrine-MpSEC22 at the NE was markedly decreased during spermiogenesis, suggesting that the composition of the nuclear envelope proteins drastically changes during spermiogenesis. Because NE-localized proteins are involved in the regulation of the nuclear shape in Arabidopsis (Goswami et al., 2020; Meier et al., 2016), the alteration in composition of the NE might be related to the nuclear reshaping during spermiogenesis in *M. polymorpha*. To our knowledge, the existence of the ER in bryophyte spermatozoids has not been reported (Renzaglia and Garbary, 2001), which infers that the ER is largely eliminated from spermatozoids. Surprisingly, however, we found that the soluble ER marker remained in the cell body of the spermatozoid (Fig. 6L). It is reported that the anterior tip of the nucleus is embedded between the spline and anterior mitochondrion during spermiogenesis in *M. polymorpha* (Kreitner, 1977a). The accumulated ER marker at the anterior region of the cell body could represent this anterior tip of the nucleus; soluble luminal ER proteins in this region might escape from degradation. Further analyses would be needed to conclusively demonstrate how the nuclear shaping and ER remodeling are coordinated during spermiogenesis.

## Material and Methods

### Plant materials and transformation

Male accession of *Marchantia polymorpha*, Takaragaike-1 (Tak-1), was used throughout this study. Plants were grown on 1/2× Gamborg’s B5 medium containing 1.4% (w/v) agar at 22 °C under continuous white light. Transformation was performed according to a previously described method (Kubota et al., 2013). Transgenic lines were selected with 10 mg·L^−1^ hygromycin B and 100 mg·L^−1^ cefotaxime for plants transformed with pMpGWB101-based binary vectors (see below), and 0.5 μM chlorsulfuron and 100 mg·L^−1^ cefotaxime for plants transformed with pMpGWB301-based binary vectors (see below). Induction of sexual organs by far-red irradiation was performed as described previously (Chiyoda et al., 2008).

### Constructions

Open reading frames (ORFs) and genomic sequences of *M. polymorpha* genes were amplified by PCR from cDNA and genomic DNA prepared from Tak-1. Amplified fragments were subcloned into the pENTR™/D-TOPO vector (Thermo Fisher Scientific) according to the manufacturer’s instructions. The primer sequences and sizes of amplified products are listed in Table S1. To construct pENTR _*pro*_Mp*DUO1* and _*pro*_Mp*IFT52*, the promoter region of each gene (5.0 and 5.4 kb, respectively) was amplified using Tak-1 genomic DNA, which was subcloned into the pENTR™/D-TOPO vector. To construct pENTR _*pro*_Mp*CEN1*, the promoter region of Mp*CEN1* (5.0 kb) was amplified with a *Sma*I site at the 3′ end and subcloned into the pENTR™/D-TOPO vector. The resultant sequence was introduced into pMpGWB307 (Ishizaki et al., 2015) using Gateway LR Clonase™ II Enzyme Mix (Thermo Fisher Scientific).

To construct pENTR _*pro*_Mp*DUO1:mCitrine*, _*pro*_Mp*CEN1:mCitrine*, and _*pro*_Mp*IFT52*:*mCitrine*, Mp*DUO1*, Mp*CEN1*, and Mp*IFT52* promoter sequences were amplified with a *Sma*I site at the 3′ end and subcloned into the pENTR™/D-TOPO vector. The cDNA of mCitrine was amplified and inserted into the *Sma*I site. To construct pMpGWB101-based and pMpGWB301-based Gateway vectors (Fig. S4D), the promoter regions followed by cDNA for mCitrine or only the promoter regions were amplified from plasmids described above, which were inserted at the *Hin*dIII site of pMpGWB101 or pMpGWB301 using the In-fusion HD Cloning System (Clontech). To construct _*pro*_Mp*DUO1*:*ST*-*Venus*, pENTR *ST*-*Venus* (Kanazawa et al., 2016; Uemura et al., 2012) was subjected to LR reaction with pMpGWB301 _*pro*_Mp*DUO1*. To construct _*pro*_Mp*DUO1: mCitrine-*Mp*SYP12A*, _*pro*_Mp*DUO1: mCitrine-*Mp*VAMP71*, _*pro*_Mp*DUO1*:*mCitrine*-Mp*GOS11*, _*pro*_Mp*DUO1*:*mCitrine*-Mp*SFT1*, and _*pro*_Mp*DUO1*:*mCitrine*-Mp*SEC22*, pENTR Mp*SYP12A*, pENTR Mp*VAMP71*, pENTR Mp*GOS11*, pENTR Mp*SFT1*, and pENTR Mp*SEC22* (Kanazawa et al., 2016) were subjected to LR recombination with pMpGWB301 _*pro*_Mp*DUO1*:*mCitrine*, respectively. To construct *_pro_*Mp*EF1*α:*SP-mCitrine-HDEL* fusion genes, pDONR _*pro*_Mp*EF1*α (Mano et al., 2018) was subjected to LR recombination with R4pMpGWB394 (Mano et al., 2018).

### Microscopy

To prepare the antheridial cells of *M. polymorpha* for observation, the antheridial receptacles between stages 3 and 5 (Higo et al., 2016) were sliced manually with a razor blade, placed on a glass slide (Matsunami), and then covered with a coverslip. To prepare the spermatids of *M. polymorpha*, antheridia were fixed for over 60 min with 4% (w/v) paraformaldehyde (PFA) in PME buffer (50 mM PIPES-NaOH, 5 mM EGTA, and 1 mM MgSO_4_, pH 6.8), and treated for 30 min with cell wall digestion buffer (1% (w/v) cellulase Onozuka RS (SERVA), 0.25% (w/v) pectolyase Y-23 (KYOWA CHEMICALPRODUCTS), 1% (w/v) BSA, 0.1% (w/v) NP-40, 1% glucose, and 1 x cOmplete™ EDTA-free protease inhibitor cocktail (Roche Applied Science) in PME buffer). The samples were placed on a glass slide and mounted with PBS containing 0.1% Hoechst 33342. To prepare the mature spermatozoids of *M. polymorpha*, collected spermatozoids were fixed for over 60 min with 4% (w/v) PFA in PBS buffer, placed on a glass slide, washed with PBS buffer three times, and mounted with PBS containing 0.1% Hoechst 33342. The prepared samples were observed under LSM780 (Carl Zeiss) with an oil immersion lens (x63).

For immunostaining of spermatids of *M. polymorpha*, we slightly modified the method described in Shimamura (2015). Antheridia were fixed for over 60 min with 4% (w/v) PFA in PME buffer and treated for 30 min with the cell wall digestion buffer. Cells were then treated with permeabilization buffer (0.01% (v/v) Triton X-100 and 1% (w/v) BSA in PME buffer) for 10 min. After washing with PME buffer three times, cells were placed on a MAS-coated glass slide, and incubated for 30 min at room temperature with blocking solution (1% (w/v) BSA in PBS buffer). After removing the blocking solution, cells were incubated with the primary antibody in PBS buffer at 4 °C overnight. After washing with PBS buffer three times, the samples were incubated for 60 min at 37 °C with the secondary antibody and 0.1% (v/v) Hoechst 33342 in PBS buffer. After washing with PBS buffer three times, slides were mounted using the ProLong Diamond Antifade reagent (Thermo Fisher Scientific). Immunostaining of mature spermatozoids was performed almost similar to that of spermatozoids, with additional centrifugation to collect spermatozoids. Samples were observed under LSM780 (Carl Zeiss) with an oil immersion lens (x63). The reactivities of the antibodies were examined using immunoblotting with cell lysates prepared from antheridia of Tak-1.

#### Antibodies

Anti α-tubulin antibody (DM1A) and anti α-tubulin antibody (B-5-1-2) were purchased from Sigma-Aldrich and used at ×1,000 dilution for immunostaining (DM1A and B-5-1-2) and immunblotting (B-5-1-2). Anti β-tubulin antibody (AA2) was purchased from Abcam and used at ×1,000 dilution for immunostaining and immunoblotting. Anti polyE antibody was purchased from AdipoGen Life Sciences (AG25B-0030-C050) and used at ×5,000 dilution for immunoblotting and ×10,000 dilution for immunostaining. Alexa Fluor 488 plus goat anti-mouse IgG (A32723) and Alexa Fluor 594 plus goat anti-rabbit IgG (A32740) were purchased from Thermo Fisher Scientific and used at ×1,000 dilution for immunostaining.

#### Gene accession numbers

Mp*SYP12A* (Mapoly0163s0015, Mp6g00050), Mp*VAMP71* (Mapoly0064s0109, Mp8g00880), Mp*SYP2* (Mapoly0187s0013, Mp8g15260), Mp*DUO1* (Mapoly0019s0071, Mp1g13010), Mp*CEN1* (Mapoly0103s0016, Mp1g00710), Mp*IFT52* (Mapoly0059s0029, Mp6g13200), Mp*GOS11* (Mapoly0001s0245, Mp1g19070), Mp*SFT1* (Mapoly0006s0109, Mp3g06390), Mp*SEC22* (Mapoly0023s0071, Mp2g11050)

## Acknowledgments

We thank Dr. Takehiko Kanazawa (NIBB) for sharing plasmids used in this study. We also thank Dr. Masaki Shimamura (Hiroshima University) for sharing the protocol for immunostaining. We thank the Model Plant Research Facility of the NIBB Bioresource Center for technical support.

## Competing interests

No competing interests declared.

## Funding

This work was supported by the following grants from the KAKENHI Program of the Japan Society for the Promotion of Science (JSPS): Grant No. 20K15824 to N.M., Grants No. 17H05850, 19H04872, and 21K06222 to K.E., and Grants No. 19H05675, 19H05670, and 21H02515 to T.U., as well as a Grant-in-Aid for JSPS fellows (to TN, Grant No. 19J13751).

## Notes

### Competing Interest Statement

The authors have declared no competing interest.

