## Supplementary Information for "Remodeling of organelles and microtubules during spermiogenesis in the liverwort *Marchantia polymorpha*" (from another version of this preprint: version 1, 2021-07-12)

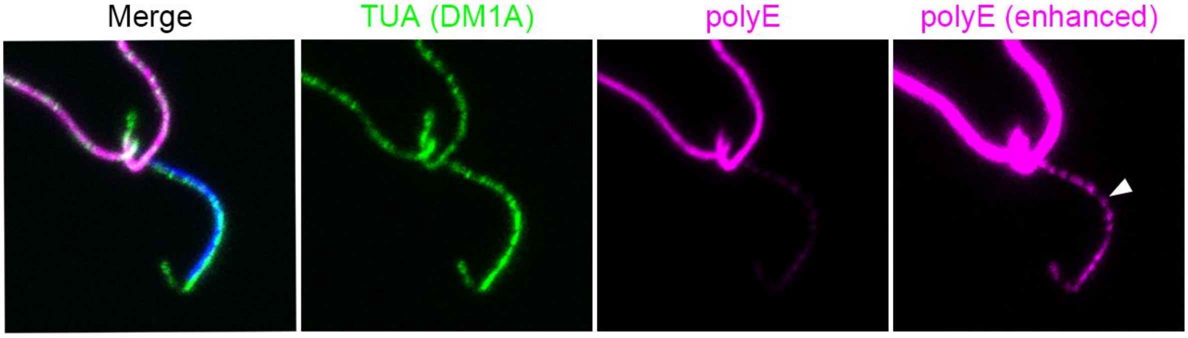


**Fig. S1. Immunostaining of spermatozoids with anti-α-tubulin and anti-polyE antibodies**

(A) Maximum intensity projection images of a spermatozoid immunostained with anti-polyE and anti-α-tubulin (TUA (DM1A)) antibodies. An arrowhead indicates the signal of polyE along the spline.

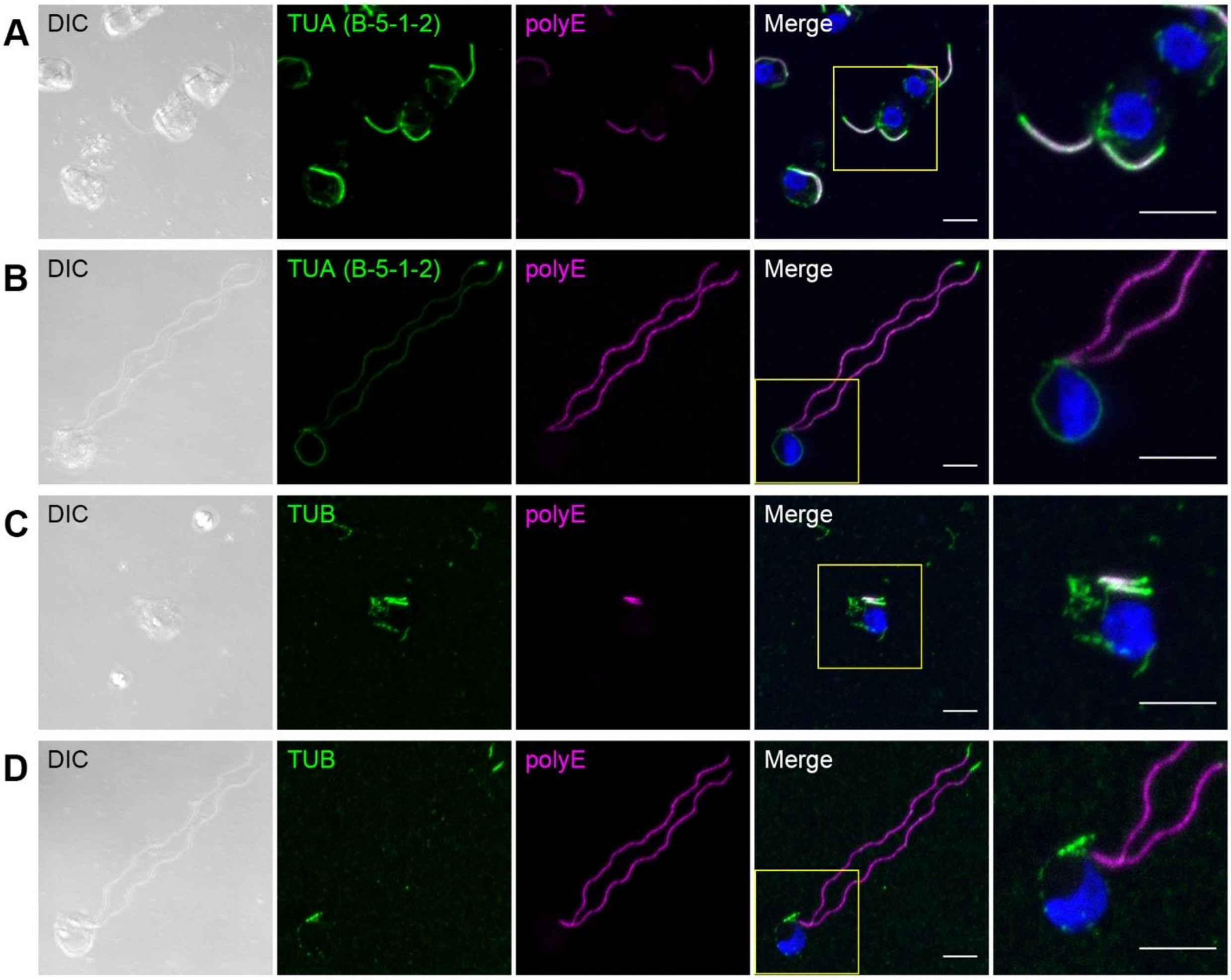


**Fig. S2. Immunostaining of spermatids with distinct anti-tubulin antibodies**

(A-D) Maximum intensity projection images of spermatids immunostained with anti-polyE and anti-α-tubulin (A and B) or β-tubulin (C and D) antibodies. The leftmost panels, differential interference contrast (DIC) images; other panels, fluorescence images. The rightmost panels are high magnification images of yellow-squared areas. Green, magenta, and blue pseudo-colors indicate fluorescence of Alexa 488 (α-tubulin or β-tubulin), Alexa 594 (polyE), and Hoechst 33342, respectively. Scale bars = 5 μm.

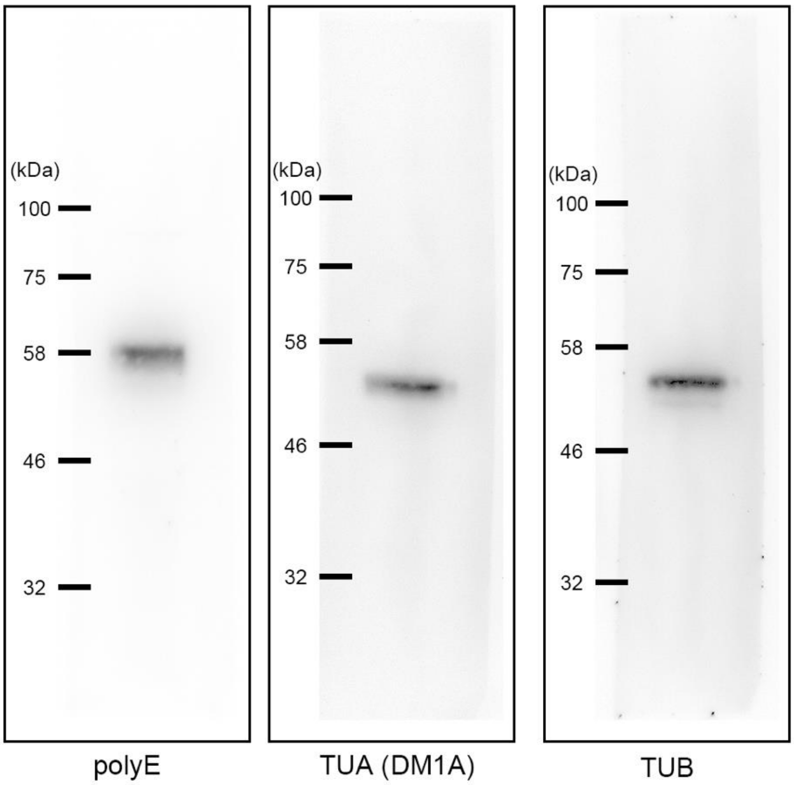


**Fig. S3. Anti-tubulin and anti-polyE antibodies react with distinct populations of tubulin.**

Immunoblotting using cell lysates prepared from antheridia of Tak-1 with anti-α-tubulin (TUA (DM1A)), anti-β-tubulin (TUB (AA2)), and anti-polyE antibodies. Black bars indicate molecular weights (kDa).

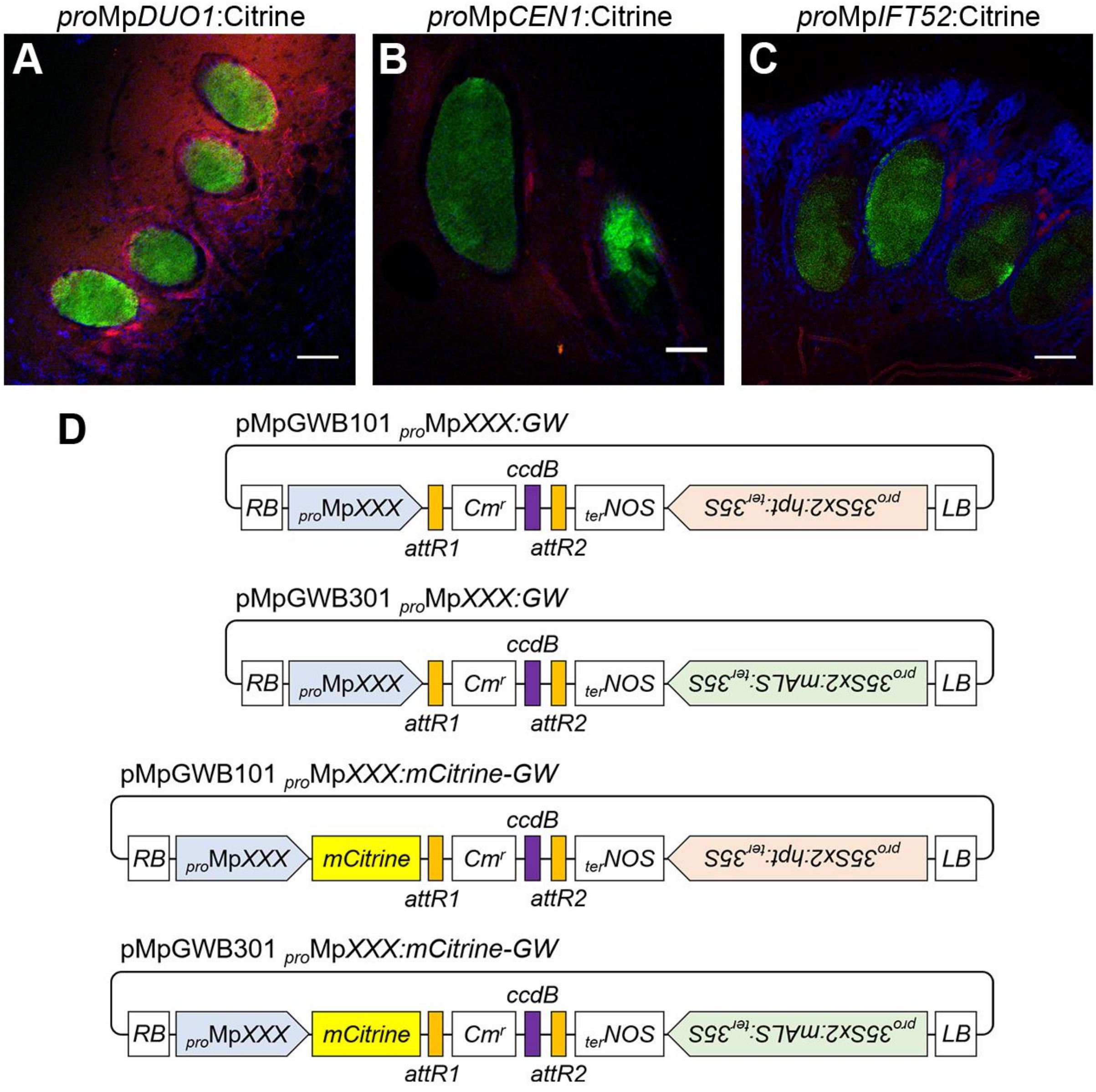


**Fig. S4. Spermatid-dominant expression by promoters of Mp*DUO1*, Mp*CEN1*, and Mp*IFT52***

(A-C) Maximum intensity projection images of sliced antheridial receptacles transformed with *pro*Mp*DUO1*:Citrine (A), *pro*Mp*CEN1*:Citrine (B), or *pro*Mp*IFT52*:Citrine (C). Green, red, and blue pseudo-colors indicate fluorescence from Citrine, anthocyanin-like compound, and chlorophyll, respectively. Scale bars = 10 μm. (D) Schematic illustration of Gateway vectors.

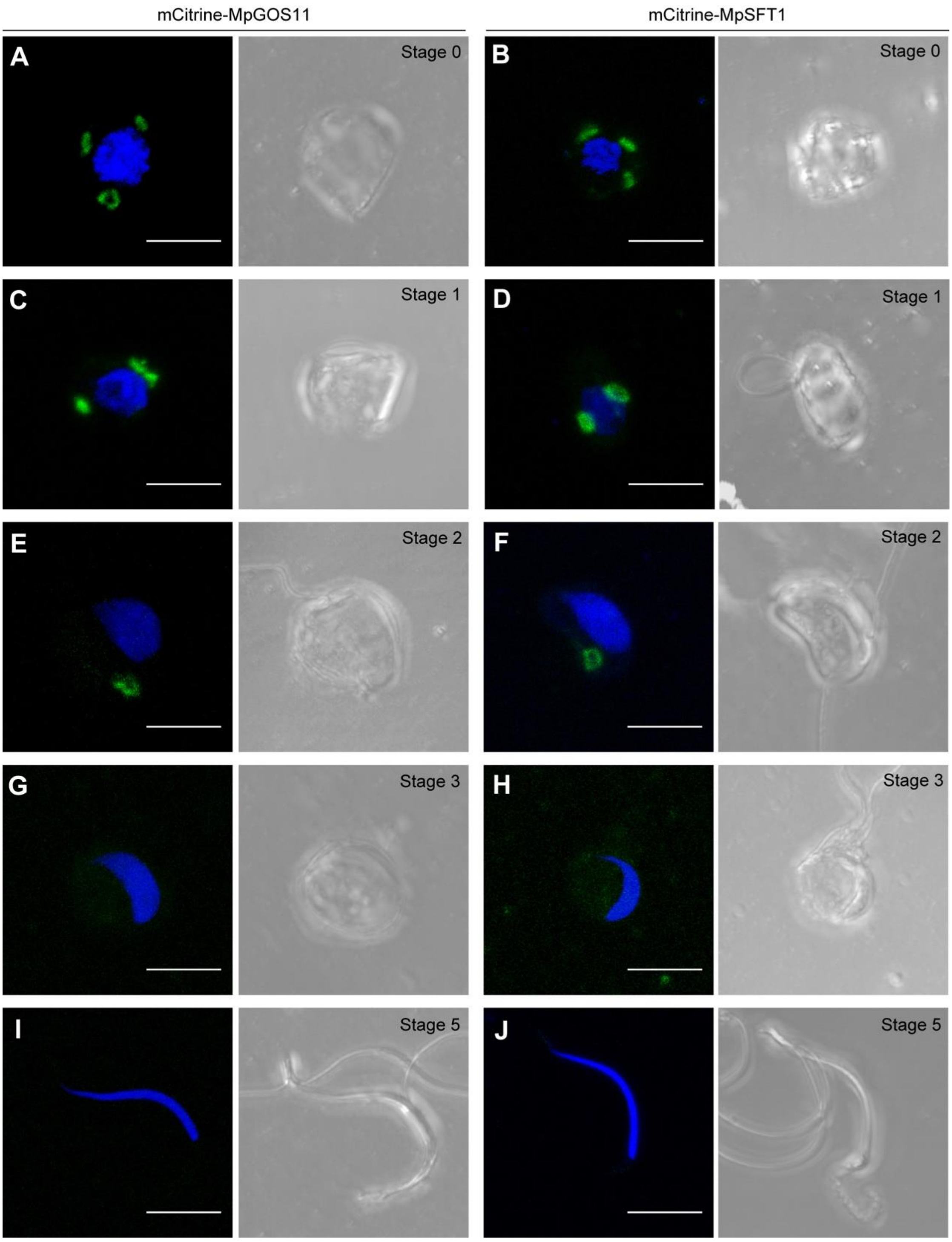


**Fig. S5. Reorganization of the Golgi apparatus during spermiogenesis**

Maximum intensity projection images of spermatids and spermatozoids expressing mCitrine-MpGOS11 (A, C, E, G, I) or mCitrine-MpSFT1 (B, D, F, H, J) driven by the Mp*DUO1* promoter. Left panels, fluorescence images; right panels, differential interference contrast (DIC) images. Green and blue pseudo colors indicate fluorescence from mCitrine and Hoechst 33342, respectively. Scale bars = 5 μm.

| primers | sequences |
|---|---|
| MpDUO1proFw | CACCGGAAAATTTTGTAGATGATTATAATTGGTATG |
| MpDUO1proRv | TTTTTATCTGAGACCAGCCACTCCG |
| MpDUO1proRv_SmaI | CCCGGGTTTTTATCTGAGACCAGCCACTCCG |
| CitrineFw_SmaI_DUO1pro | CTCAGATAAAAACCCATGGTGAGCAAGGGCGAGGA |
| Citrine-stopRv_SmaI | GCGCCCACCCTTCCCGCTTCCTCCTCCCTTGTACAGCTCGTCCATGC |
| MpDUO1proFw_HindIII | GGCCAGTGCCAAGCTGGAAAATTTTGTAGATGATTATAATTGGTATG |
| MpDUO1proRv_HindIII | TTTGTACAAACTTGTTTTTATCTGAGACCAGCCACTCCG |
| CitrineRv_HindIII | TTTGTACAAACTTGTCTTGTACAGCTCGTCCATGCCG |
| MpIFT52proFw | CACCGGTTAACATCCGATAGATTTGTGGAAG |
| MpIFT52proRv | CTCTTAGAATGGATGCGCATGC |
| MpIFT52proRv_SmaI | CCCGGGCTCTTAGAATGGATGCGCATGC |
| CitrineFw_SmaI_IFT52pro | CCATTCTAAGAGCCCATGGTGAGCAAGGGCGAGGA |
| MpIFT52proFw_HindIII | GGCCAGTGCCAAGCTGGTTAACATCCGATAGATTTGTGGAAG |
| MpIFT52proRv_HindIII | TTTGTACAAACTTGTCTCTTAGAATGGATGCGCATGC |
| MpCEN1proFw | CACCACCAGATGAAACCGAGTCCGTGAG |
| MpCEN1proRv_SmaI | CCCGGGGGTGGAATCGAACGATGACACAGG |
| CitrineFw_SmaI_CEN1pro | TTCGATTCCACCCCCATGGTGAGCAAGGGCGAGGA |
| MpCEN1proFw_HindIII | GGCCAGTGCCAAGCTACCAGATGAAACCGAGTCCGTGAG |
| MpCEN1proRv_HindIII | TTTGTACAAACTTGTGGTGGAATCGAACGATGACACA |

**Table. S1. List of primers used in this study**
